# Durotaxis of passive nanoparticles on elastic membranes

**DOI:** 10.1101/2021.04.01.438065

**Authors:** Ivan Palaia, Alexandru Paraschiv, Vincent Debets, Cornelis Storm, Anđela Šarić

## Abstract

The transport of macromolecules and nanoscopic particles to a target cellular site is a crucial aspect in many physiological processes. This directional motion is generally controlled via active mechanical and chemical processes. Here we show, by means of molecular dynamics simulations and an analytical theory, that completely passive nanoparticles can exhibit directional motion when embedded in non-uniform mechanical environments. Specifically, we study the motion of a passive nanoparticle adhering to a mechanically non-uniform elastic membrane. We observe a non-monotonic affinity of the particle to the membrane as a function of the membrane’s rigidity, which results in the particle transport. This transport can be both up or down the rigidity gradient, depending on the absolute values of the rigidities that the gradient spans across. We conclude that rigidity gradients can be used to direct average motion of passive macromolecules and nanoparticles on deformable membranes, resulting in the preferential accumulation of the macromolecules in regions of certain mechanical properties.

## Introduction

The targeted transport of nano-objects is of great interest for numerous applications, ranging from engineering novel nanomaterials to designing efficient strategies for the delivery of nanoparticles. A multitude of approaches can be employed to manipulate the transport of nanoscopic objects to specific destinations in living and synthetic matter. For example, cells use motor proteins to transport macromolecules and organelles across the cytoplasm [1, 2]. Viruses use cellular pH gradients for cellular entry and uncoating [3]. In a synthetic context, electric, thermal, and chemical gradients can be used to artificially direct the motion of molecular cargoes within nanochannels [4–7] or on surfaces [8].

A commonly observed method to guide transport involves the usage of rigidity gradients. Perhaps the best known example of such transport is *durotaxis*, which was first observed in cells and was defined as the tendency of living cells to migrate towards regions of higher stiffness [9]. Existing physical models attribute this phenomenon either to rigidity-dependent persistence of motion, implying that cells sense and adapt to the absolute rigidity of the underlying substrate, or to gradient-dependent forces, implying that cells sense rigidity gradients on the scale of a single cell [10–12]. In both cases – in the former more subtly than in the latter – energy is implied to be required to drive directed motion.

Rigidity-guided migration is, however, not only restricted to active systems, but has also been observed in a range of passive nanoscopic scenarios. Gradients in the rigidity or in the strain field of a substrate have been used to orient motion of graphene nanosheets [13] and nanoflakes [14]. Water droplets can undergo reverse durotaxis, migrating towards softer regions of a surface to increase wetting [15], whereas non-wetting droplets undergo regular durotaxis for the opposite reason [16]. Analogously, polymer droplets tend to migrate to stiffer surfaces, where their Van der Waals energy is minimised [17]. All these passive systems share a similarity: the nano-object migrates as to increase its contact with the underlying substrate and lower the system’s potential energy.

In this paper, we study the durotactic motion of passive nanoscopic objects diffusing on deformable membranes. Biological membranes have a diverse composition and contain different species of phospholipids and proteins [18], whose expression on the cell surface can exhibit spatio-temporal dependence [19, 20]. This can possibly result in non-uniform mechanical properties across its surface [21, 22]. Such *lateral* stiffness heterogeneities have been observed *in vivo* and *in vitro*, for instance in protozoa [23, 24], red blood cells [25, 26], T-cells [27], rat neurons and HeLa cells [28]. In these systems, the variation in the elastic moduli (which may differ by as much as a factor of 15 [23]) is thought to be functional for the cell. Heterogeneity has been attributed to the chemical composition of the membrane, to the mechanics of the underlying cytoskeleton, and to their mutual interplay [29–32]. Recent computational work has reported that a nanoparticle bound to a microphase-separated multi-component membrane exhibits preference for a given phase, as a consequence of different bending rigidities between the two phases [33]; also, membrane elasticity has been proposed to influence the persistence of motion of a nanoparticle that actively cleaves the underlying substrate via the so-called burnt bridge mechanism [34].

The key idea in the present paper is that stiffness inhomogeneity alone, even in the absence of any active mechanism, might direct the dynamics of macromolecules bound to the membrane and lead to their preferential accumulation in regions of optimal rigidity. The aim of this paper is to understand the physics of this effect from first principles and to propose it as a novel sorting mechanism in its own right. We use molecular dynamics simulations to study the preferential localisation of a hard spherical nanoparticle on a non-homogeneous elastic membrane. We place an adhering nanoparticle on a fluctuating membrane divided in two halves of different bending rigidities, as shown in Fig. 1. We show that a difference in the values of the bending rigidity between the two regions of the membrane is sufficient to drive the nanoparticle’s localisation to one side. We provide a theoretical underpinning for this phenomenon, based on free energy calculations and analytical estimates. Depending on the absolute values of the two rigidities, we observe motion both up and down the rigidity gradient, thereby effectively demonstrating both regular and negative durotaxis.

**FIG. 1:**
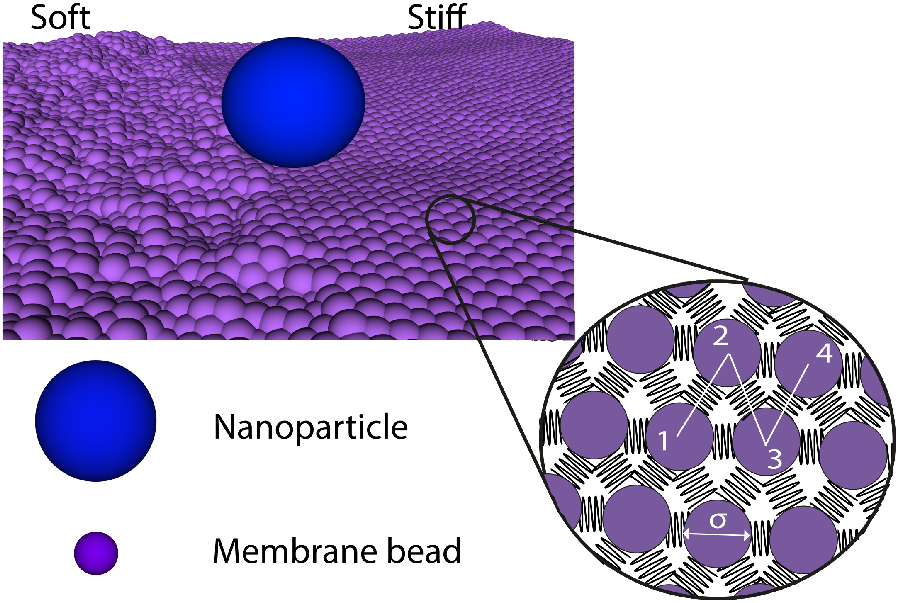
Illustration of a hard spherical nanoparticle on an elastic membrane that contains two halves of different rigidities. The right half has a greater bending rigidity than the left half. The membrane is described as a triangulated elastic surface, where the bending rigidity is controlled by the dihedral potential between adjacent triangles (1-2-3 and 2-3-4). The diameter of the nanoparticle is *σ*NP = 10 *σ*, where *σ* is the simulation unit of length and the diameter of one membrane bead. See Methods for details.

## Results and Discussion

### Adhesion non-monotonically depends on the bending rigidity

To investigate how the substrate’s stiffness influences the preferential localisation of a nanoparticle, we first placed the nanoparticle on uniform membranes of different rigidities, *K*_*b*_, varied between *K*_*b*_ = 0.01 *k*_B_*T* and *K*_*b*_ = 500 *k*_B_*T*. For every set of simulations, we tracked the average nanoparticle coordination number, defined as the number of membrane particles within range of interaction with the nanoparticle, as well as the total adhesion interaction energy between the membrane and nanoparticle.

The nanoparticle’s adherence to the membrane is influenced by the balance of three terms: the energetic cost of locally deforming the membrane, the energy gain due to the adhesion of the nanoparticle, and an entropic effect related to membrane fluctuations. Depending on the value of the bending rigidity, the contact between the nanoparticle and the membrane exhibits four distinct regimes as shown in Fig. 2. For low values of the bending rigidity, the membrane is conspicuously corrugated and its fluctuations hinder full contact with the particle (regime I). As the rigidity is increased, the membrane fluctuations are gradually suppressed and the nanoparticle can be wrapped more by the membrane, leading to a substantial increase in the coordination number and in the absolute value of the total adhesion energy (regime II). At values of *K*_*b*_ above ∼ 2 *k*_B_*T*, the mechanical cost of deforming the membrane overcomes the adhesion term, resulting in decreased adherence (regime III). The entropic term can be neglected at high rigidities. For very high values of the bending rigidity, the local membrane deformation imposed by the nanoparticle is prohibitively costly and the coordination number eventually saturates at a low value (regime IV). These results show that there is an optimal value of the bending rigidity that maximises the adherence of the nanoparticle diffusing on the surface of a fluctuating membrane. We expect this non-monotonic behaviour of the adhesion interaction to have a direct impact on the preferential localisation of the nanoparticle on the membrane.

**FIG. 2:**
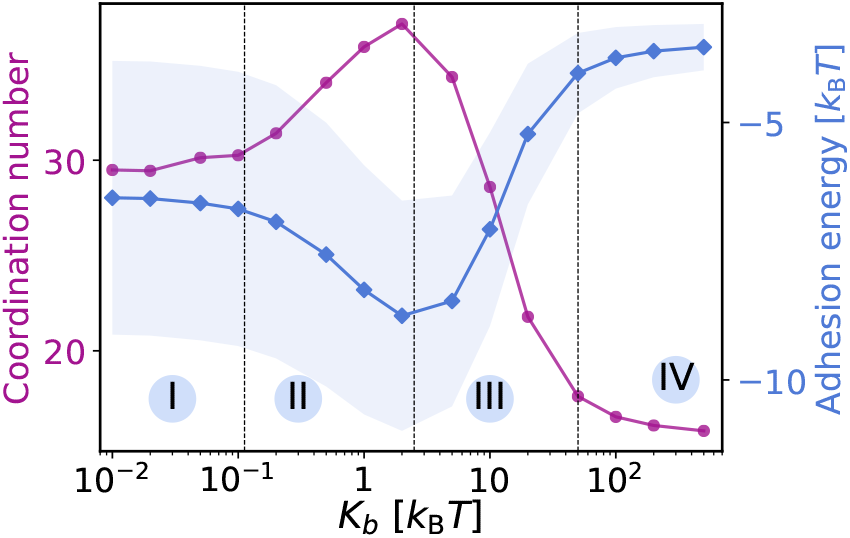
Nanoparticle’s time-averaged coordination number and adhesion energy as a function of bending rigidity of a uniform membrane. The nanoparticle contact with the membrane follows a non-monotonic pattern, with a maximum at a value of *K*_*b*_ *∼* 2 *k*_B_*T*. Four different regimes appear, described in the text. The shaded area indicates the standard deviation of the adhesion energy distribution.

### Nanoparticle localisation is strongly influenced by the stiffness gradient

According to the previous observations, a nanoparticle adsorbed on an inhomogeneous membrane should preferentially migrate to regions where the adhesion is maximised in order to minimise the system’s total energy, thus resulting in a form of durotaxis. We test this hypothesis by placing the nanoparticle on a membrane divided in two halves of different rigidities (as illustrated in Fig. 1). We expect that the local bending rigidity will influence the statistical partitioning of the particle between the two membrane regions. Fig. 3 illustrates how the preference of the nanoparticle for either side of the membrane changes depending on the rigidity of each surface. Here, the nanoparticle localisation is quantified by counting the time spent by the particle on each side of the membrane. The nanoparticle shows substantial preference for one side of the membrane.

**FIG. 3:**
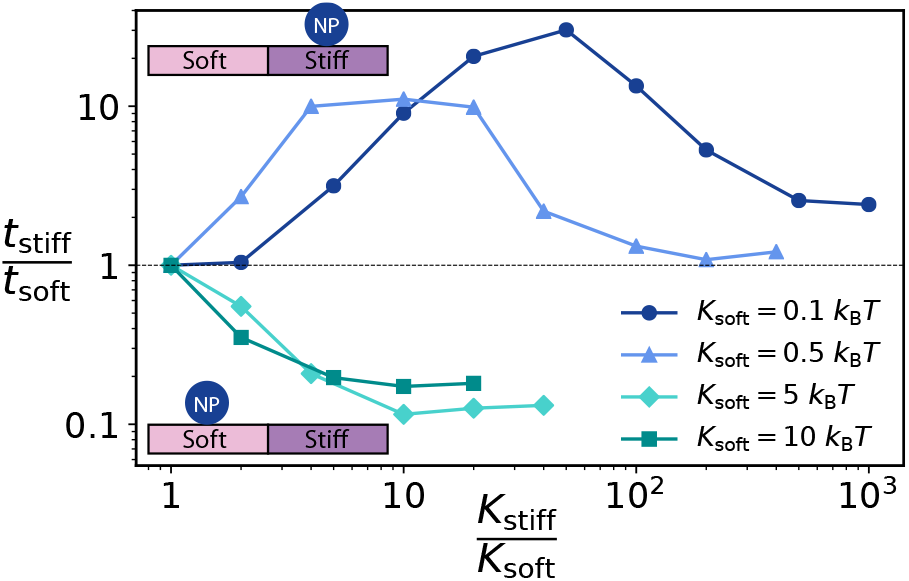
Partitioning of the nanoparticle between the rigid and the soft region of the membrane. *t*_soft_ and *t*_stiff_ are the amounts of time spent by the particle on each of the two surfaces, of rigidity *K*_soft_ and *K*_stiff_ respectively. As shown by the drawings, values of *t*_stiff_ */t*_soft_ above 1 indicate preferential affinity for the stiff side, and vice versa. For each curve, *K*_soft_ is fixed (see legend).

In particular, we distinguish two different durotactic regimes, depending on the bending rigidity *K*_soft_ of the softer region. For small *K*_soft_, the entropic effects seem to dominate over the energy ones and the nanoparticle preferentially localises on the rigid side (top half of Fig. 3). In this case, the particle follows a proper durotactic motion, displaying a tendency to migrate towards stiffer regions. Conversely, if *K*_soft_ is larger, the preference will be reversed (bottom half of Fig. 3). These findings are in agreement with the variation of the nanoparticle’s average adherence shown in Fig. 2. The particle will have greater preference for states that maximise its contact with the membrane. As such, nanoparticles will exhibit a tendency to migrate to regions of rigidity closer to the value of maximum adherence (∼ 2 *k*_B_*T*) observed in Fig. 2. The non-monotonic behaviour of the curves in the top half of Fig. 3 (proper durotactic regime) also qualitatively follows the trend in adherence: the partitioning shifts slightly in favour of the softer region for large *K*_stiff_, as wrapping favours very soft membranes compared to very stiff (Fig. 2). Altogether, these results demonstrate that adherence greatly influences the particle’s motion on an inhomogeneous membrane.

### Free energy analysis

To better understand how adherence (Fig. 2) and, more importantly, durotaxis (Fig. 3) depend on the bending rigidity of the membrane, we outline a simple analytical model to study the free energy of adhesion. This free energy can be directly compared to our earlier results, since the ratio of time spent on either side of the membrane should be proportional to the exponential of the free energy difference between the stiff-bound and soft-bound state. In other words, given any two values of rigidity *κ*_soft_ and *κ*_stiff_ [53], the nanoparticle partitioning will depend only on Δ*F*_soft*→*stiff_ = *F* (*κ*_stiff_) − *F* (*κ*_soft_). The key observation is that we can choose as a reference state to measure free energies the one in which the particle is unbound from either half of the membrane: we call *F* (*κ*) the free energy the system gains if a previously detached nanoparticle is absorbed onto a membrane of rigidity *κ*, with *κ* = *κ*_stiff_ or *κ*_soft_.

The approach we use to compute the free energy is the following (see Methods for mathematical details). We aim at writing a constrained free energy, depending explicitly on the amount of adhered surface *A*; the minimum of this constrained free energy will then give the optimal wrapping and the equilibrium free energy value. Now, *F* includes a bending term *E*_bending_ ∝ *κA*, an adhesion term *E*_adhesion_ ∝ −*A*, and an entropic term −*TS*. A fourth term, related to membrane tension, could be inserted; however, the simulated membrane is effectively tensionless and can only sustain local transient stretching, so we neglect surface tension in our thermodynamic treatment. For simplicity, we restrict ourselves to the limit of strong adhesion. While the energetic terms *E*_bending_ and *E*_adhesion_ are easily written down in a coarse grained fashion neglecting fluctuations (Methods Eqs. (5) and (6)), a correct evaluation of the entropy of a generic confined membrane is a more challenging undertaking [35–37]. We assume that bound membrane beads, which are strongly confined close to the nanoparticle surface, only fluctuate along the confinement axis. This allows us to estimate the amplitude *δh* of these fluctuations by invoking the equipartition theorem and using a quadratic expansion of the adhesion potential around its minimum. We notice that adhesion strongly suppresses fluctuations: in particular, short-wavelength modes tend to be controlled by bending, both when the nanoparticle is bound to the membrane and in the unbound reference state; on the contrary, long-wavelength modes are suppressed in the bound state more than in the reference state, due to adhesion. The entropy difference is then computed within the approximation that beads fluctuate independently from each other and perpendicularly to the surface. This gives a 1D-ideal-gas-like entropy contribution, that is proportional to the number of bound beads (i.e. to the adhered surface *A*) and depends on the ratio of bending rigidity versus adhesion energy (Methods Eqs. (8), (12) and (14)).

Interestingly, this simplified model can reproduce the results of our simulations and help us make sense of the entropic effects. In Fig. 4 we show the free energy resulting from the minimization process, broken down in its three components. For low bending rigidity values (I), entropy limits wrapping, as observed in simulations (Fig. 2), and counterbalances adhesion energy. As rigidity increases (II), if wrapping stays minimal, the fluctuations of the reference unbound state decrease, until they have the same amplitude as in the bound state and the entropy loss due to adhesion vanishes. As a result the system can afford a larger adhered surface *A*. As *A* gets larger, though, new bound large-wavelength modes become available, which are suppressed by adhesion more than they would be in the unbound state. This yields a new source of entropy loss, which limits the growth in the adhered surface and sets the optimal wrapping. Increasing *κ* further (III), the bending energy per unit surface becomes comparable with (or larger than) the adhesion energy per unit surface and bending becomes more and more unfavourable. At the same time, entropy loses relevance. This is because the cutoff between adhesion-limited and bending-limited modes gets shifted to wavelengths that become unphysically larger than 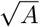, so that all available modes are now bending-limited. Consequently, the entropy difference *S* between a bound and an unbound membrane gradually goes to zero. In this regime, optimal wrapping is dominated by a competition between bending energy and adhesion energy and can only decrease with *κ*, until the point (IV) where *A* becomes as small as possible (virtually just one point of nanoparticle-membrane contact).

**FIG. 4:**
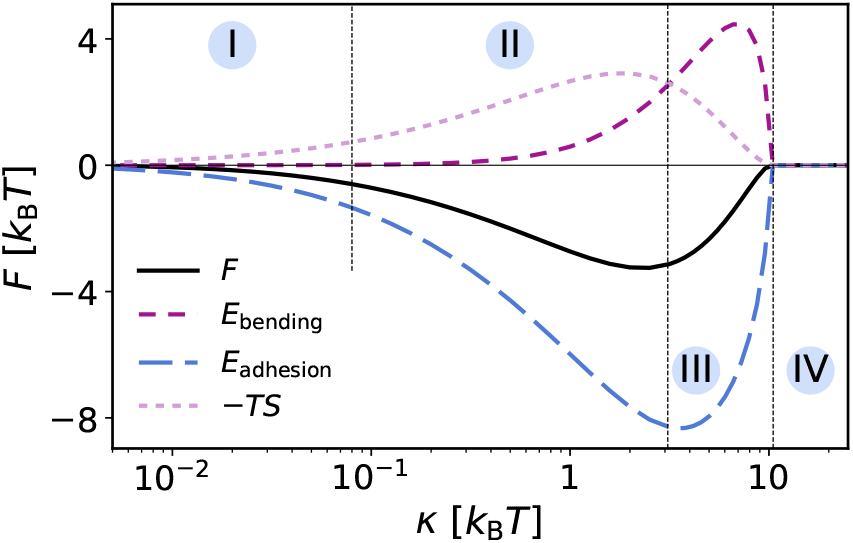
Free energy *F* as a function of bending rigidity *κ*, within the analytical approximation. Dashed and dotted lines correspond to bending energy *E*_bending_, adhesion energy *E*_adhesion_ (representing also the wrapped area *A*), and entropic term − *T S* (where *S* is entropy). The sum of these three terms is the total free energy *F*. The adhesion energy, which is directly proportional to the adhered surface *A*, reproduces the trend from Fig. 2. See Methods for details.

This analysis demonstrates that the free energy curve (thus including entropic effects) follows the same qualitative trend as the adhesion energy curve, clarifying why the durotactic behaviour observed in Fig. 3 can be explained with the nanoparticle’s preference for the higher membrane wrapping (Fig. 2). In addition, since 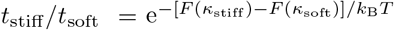, the logarithmic partitioning curves shown for given *K*_soft_ in Fig. 3 probe precisely *F* (*κ*). In particular, for each curve, a different *κ*_soft_ (or *K*_soft_) is taken as zero-energy reference, so that each curve represents a different part of *F* from Fig. 4.

The shape of the free energy from Fig. 4 is robust against changes in the size *R* of the particle, with its minimum point *κ*_min_ growing roughly as *R*^2^ and its minimum value *F* (*κ*_min_) as *R*. Similarly, increasing adhesion deepens and shifts the minimum to the right, presumably up to a point (never reached in our simulations) where the particle is fully wrapped.

### Mechanical mechanisms and kinetics

To provide an additional comparison with our theory, we measure the potential of mean force for a membrane-bound nanoparticle crossing the interface between two regions of different rigidities, by means of umbrella sampling (see Methods). Adopting similar settings as in Fig. 3, we fix the rigidity of the softer side and vary that of the stiffer side. In the case of low *K*_soft_ = 0.5 *k*_B_*T*, the free energy decreases significantly when passing to the stiff side of the membrane (Fig. 5a). This finding is consistent with the trend observed for the same *K*_soft_ in Fig. 3. The free energy difference Δ*F*_soft*→*stiff_ corresponds to the difference in potential of mean force between positive and negative *x*, far from the interface. It exhibits non-monotonic behaviour that matches the one observed in Fig. 4, displaying a deep minimum at intermediate values of bending rigidity *K*_stiff_ 10 ≃ *k*_B_*T*. Interestingly, the free energy difference favours the stiff side even at very large *K*_stiff_, as there is some non-zero adhesion present at the particle-membrane contact even in that case: this indicates a limit of the analytical model, whose continuous treatment of adhesion assumes zero energy gain at infinite rigidity. In the case of a larger *K*_soft_ = 5 *k*_B_*T* (Fig. 5b) the free energy increases in value when passing from the soft to the stiff side, again in agreement with the corresponding curve of Fig. 3.

**FIG. 5:**
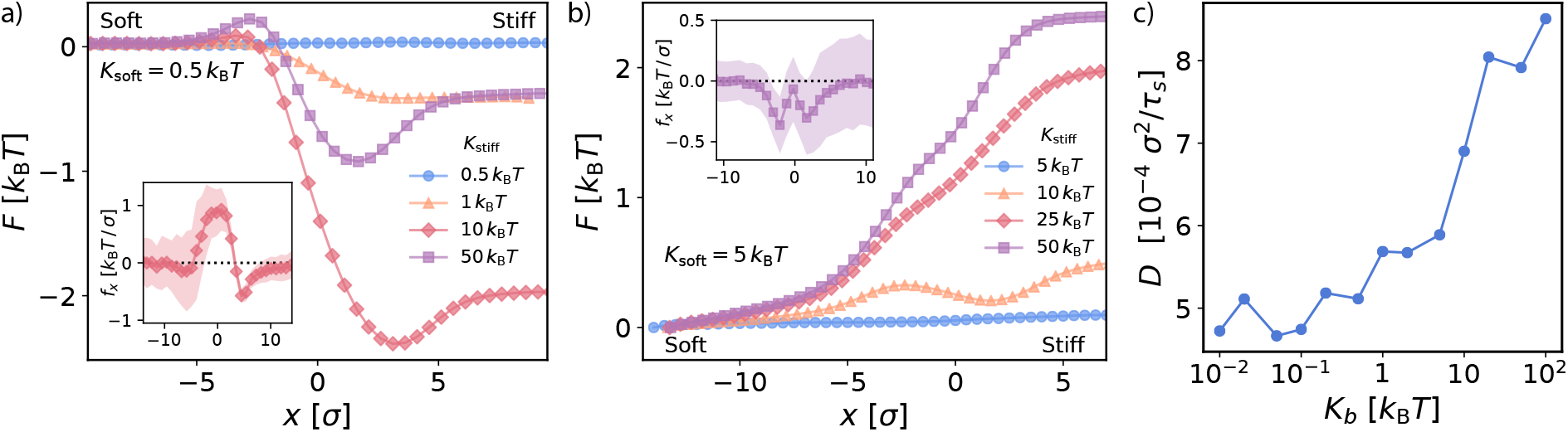
Durotaxis is caused by a net force acting at the interface, not by rigidity-dependent kinetics. (a) and (b) Potential of mean force (free energy) across the direction of the stiffness gradient (*x*). The softer region (left-hand side) has bending rigidity *K*_soft_ = 0.5 *k*_B_*T* (a) and *K*_soft_ = 5 *k*_B_*T* (b). The insets show the average force *fx* acting on the nanoparticle: *fx >* 0 means that the particle is pushed toward positive *x* values, and vice versa. The bending rigidities of the rigid side are in the legend. (c) Diffusion coefficient of the nanoparticle, moving without constraints on a uniform membrane, projected on the *xy* plane (parallel to the average membrane position).

Figs. 5a-b show that the change in free energy occurs on a length scale comparable to the diameter of the nanoparticle (10 *σ*). More explicitly, their insets represent the spatial distribution of the average force *f*_*x*_ acting on the nanoparticle along the direction of the gradient *x*. These plots show that the particle experiences a pulling force in the direction of lower free energy, or maximum wrapping, every time it approaches the rigidity gradient. This force is the microscopic mechanism behind the passive durotaxis we observe, falling in the category of gradient-sensing mechanisms. These have been shown – albeit in the different context of cell durotaxis [12] – to be more efficient than absolute-rigidity-sensing mechanisms, also called durokinetic because they only rely on the fact that kinetic parameters depend on the local rigidity [38]. To assess the potential copresence of a durokinetic process, we computed the projected 2D nanoparticle diffusion coefficient *D* (Fig. 5c). Durokinesis predicts motion in the direction of larger diffusivity, which is roughly proportional to the persistence time: we would therefore expect *D* to first grow and then decrease with stiffness. This is not the case; besides, its increase is limited to a factor *<* 2, much smaller than in related studies [12, 38]. We conclude that kinetics cannot be the dominant durotactic mechanism here, in agreement with thermodynamic considerations detailed in the Conclusions.

The rigidity dependence of the projected diffusion coefficient is however interesting in its own right [39]. It does not trivially follow wrapping; instead, it decreases monotonically starting from its free-particle value, as rigidity decreases. This is qualitatively consistent with purely geometric effects attributed to surface corrugation [40, 41].

### Durotaxis of multiple nanoparticles

The observed single-particle durotactic behaviour can lead to collective migration of multiple membrane-adsorbed particles. However, membrane-mediated interactions between multiple particles can render the resulting behaviour more complex. As a proof of principle, we explore the collective preferential migration of an ensemble of nanoparticles adhered on the inhomogeneous membrane. To do so, we place 36 identical particles uniformly on the membrane surface (corresponding to a projected surface packing fraction *ϕ* ≃ 0.28) and we measure partitioning. The particles are allowed to diffuse freely across the membrane regions and they interact with each other through volume exclusion. As shown in Fig. 6a, multiple particles behave in a manner analogous to what was observed for a single particle (Fig. 3). For low soft-side rigidity, they prefer transferring to the rigid side (first two bars in Fig. 6a); in case of similar values of bending rigidity, no significant preference for either surface is observed (third bar); for large soft-side rigidity, particles migrate to the softer side (fourth bar). These observations suggest that collective transport of macromolecules can be achieved solely through gradients in the local environment’s bending rigidity.

**FIG. 6:**
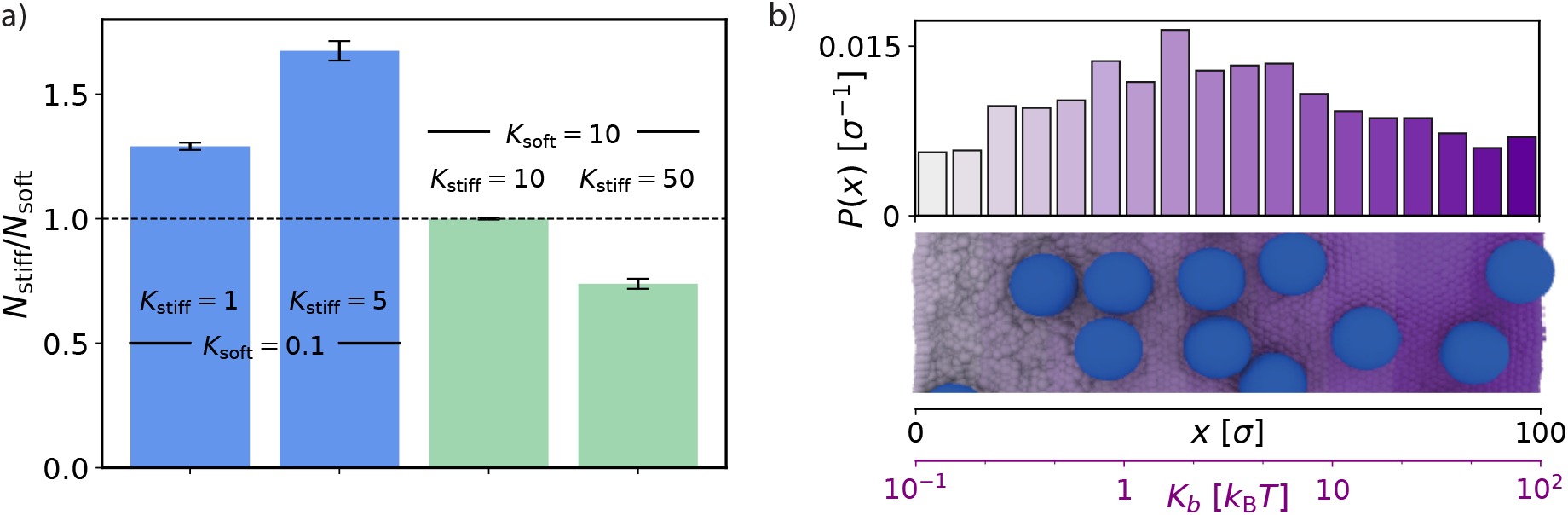
Partitioning of a collection of particles. a) Ratio between average number of particles on the stiff and on the soft side. The considered bending rigidities are specified on each bar. The results are averaged over 100 simulation repeats. b) Probability distribution of nanoparticles on a membrane in which rigidity is increased step-wise along *x*, from one end to another, in an exponential manner. A section of a simulation snapshot highlights rigidity increments on the membrane and shows accumulation of particles in its central region.

To further investigate whether more intricate rigidity gradients can induce a certain degree of localisation on the membrane, we engineer a membrane in which the bending rigidity increases gradually, logarithmically, from *K*_*b*_ = 0.1 *k*_B_*T* at one end to *K*_*b*_ = 100 *k*_B_*T* at the other. The nanoparticles statistically accumulate in the central region (Fig. 6b) which coincides with the values at which free energy in minimized (Figs. 4 and 5), or, equivalently, average adhesion is maximised (Fig. 2).

This finding suggests that gradients in membrane rigidity are sufficient to bias local concentration of particles on soft membranes. We propose this as a relevant mechanism to sort proteins on membranes or to guide diffusion of macromolecules to their target locations during physiological processes, or in artificial setups.

## Conclusions

We demonstrated that rigidity gradients can provide an intrinsic driving force for guiding the motion of passive spherical nanoparticles on deformable non-uniform membranes. This behaviour originates purely from the minimisation of the system’s free energy, causing a net force where the rigidity is non-uniform: it is not a consequence of a rigidity-dependent persistence of motion, as was proposed for cellular durokinesis [38]. Indeed, durokinesis concerns non-equilibrium systems, whose activity is hidden in the stochastic equations of motion (see [42] for a general discussion). Our system is instead intrinsically passive and must behave according to standard statistical mechanics, irrespective of the kinetics. This mechanism leads to more efficient transport [12], realised by gradient sensing on the nanoparticle scale. The fact that motion follows the direction of maximum wrapping carries analogies with cell durotaxis, where in some cases stronger adhesion is observed on stiffer substrates [11].

We observed a non-monotonic dependence in the particle’s average adhesion to the membrane. In particular, in the regime of most biologically relevant bending rigidity (∼ 20 *k*_*B*_*T*), we found that the particle migrates toward softer surfaces, contrary to what is expected for durotaxis observed in cellular systems.

This non-monotonic dependence was explained by the competition between three terms: the entropic effects which are relevant on very soft membranes, the energetic gain from adhesion, and mechanical effects due to membrane bending which penalise local deformations. A deeper understanding of our findings was provided by free energy calculations, both in simulations and with the help of an analytical theory. In particular, we proposed a simple model to account for the entropic term, whose dependence on the bending rigidity is highly nontrivial and incorporates two competing effects. On the one hand, increasing rigidity decreases the entropy lost upon adhesion, because it reduces fluctuations in the unconstrained membrane. On the other hand, for this reason, a higher rigidity allows for an increase in wrapping, which in turn unlocks longer-wavelength adhesion-limited modes and increases the entropy difference again. This feedback mechanism controls the free energy.

Finally, we showed that gradients in rigidity are enough to drive spontaneous oriented motion of many particles. This phenomenon might serve as a method to direct macromolecules toward specific functional sites on the cellular surface. Rigidity gradients have recently been shown to affect phase separation in a 3D elastic medium, comparable to the cytoskeleton [43, 44]: our results suggest that they might also act as a passive particle-sorting mechanism in a 2D environment. In particular, given the high sensitivity of the free energy with respect to particle size – it scales as the square of the particle radius – the mechanism might prove especially efficient in segregating single proteins from dimers or larger aggregates, serving a purpose similar to phase separation phenomena. Moreover, these findings could have potential impact on the development of nanodevices that involve the transport of molecular cargoes to a targeted region, for instance in the development of artificial drug delivery systems.

## Acknowledgements

We acknowledge support from the Engineering and Physical Sciences Research Council (A.P. and A.Š.), the Royal Society (A.Š.) and the European Research Council (I.P. and A.Š.).

## Methods

### A. Simulation details

The simulated system consists of a colloidal nanoparticle placed on a fluctuating elastic membrane, as shown in Fig. 1. The membrane contains 7832 beads placed on the nodes of a triangulated hexagonal mesh. The membrane beads and nanoparticle have a diameter *σ* and *σ*_NP_ = 10 *σ* respectively. The membrane beads are interconnected via harmonic springs, obeying to the following potential:

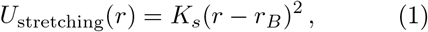

with stretching constant *K*_*s*_ = 18 *k*_B_*T* /*σ*^2^ and equilibrium bond length *r*_*B*_ = 1.23 *σ*. The membrane bending rigidity is controlled by the following dihedral potential between the opposite vertices of triangles sharing an edge (Fig. 1):

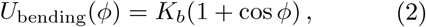

where *ϕ* is the corresponding dihedral angle, and *K*_*b*_ is the harmonic constant that controls the membrane’s bending rigidity. In our simulations, we explore a wide range of values of *K*_*b*_, between 0.01 *k*_B_*T* and 500 *k*_B_*T*.

Besides the stretching and bending terms, membrane beads interact with each other via a repulsive Weeks-Chandler-Andersen (WCA) potential [45] to impose self-avoidance:

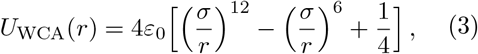

with a radial cutoff of *r*_*c*_ = 2^1*/*6^ *σ. ε*_0_ is set to 5 *k*_B_*T*.

The nanoparticle interacts with the membrane via a truncated-shifted Lennard-Jones potential:

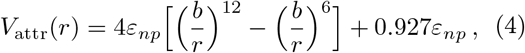

where *b* = 5.5 *σ* and *ε*_*np*_ = 10 *k*_B_*T*. A radial cutoff of *r*_*c*_ = 6.5 *σ* is used. The parameters were finely tuned to prevent the full wrapping and subsequent engulfment of the nanoparticle and to allow it to diffuse laterally on the membrane surface.

The simulation box is initialised with sides of length *L*_*x*_ = *L*_*y*_ = 100 *σ* and *L*_*z*_ = 40 *σ*. Periodic boundary conditions are applied in the *x* and *y* direction and the box is kept fixed in the *z* direction. In simulations involving a heterogeneous membrane (Figs. 1, 3, 6, and insets of 5a-b), Lennard-Jones walls with potential depth *ε* = 10 *k*_B_*T* confine the nanoparticle to the simulation box, preventing it from crossing the periodic boundaries. These walls do not interact with the membrane beads. To compute partitioning (Fig. 3), we consider time spent in the region of space more than ~ 10 *σ* away from the soft-stiff interface and more than ∼ 10 *σ* away from the closer *x* wall, in order to reduce boundary effects (this region is 30 *σ* long on either side of the membrane). In simulations involving a homogeneous membrane (Figs. 2 and 5c), the walls only act in the *z* direction, so the nanoparticle can diffuse in the *xy* plane to neighbouring image cells. Unwrapped trajectories are then used to compute mean square displacements and diffusion coefficients.

The simulations were run in the isoenthalpic-isobaric (NPH) ensemble, with a zero lateral pressure and at constant temperature. In order to replicate the stochastic dynamics of the real system, we used a Langevin thermostat with friction coefficient *γ* = 1 *m/τ*_0_, where *m* is the particle mass (equal to *m*_0_ for membrane beads and to 100 *m*_0_ for the nanoparticle) and 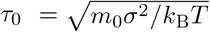 the natural time unit of the simulation. The simulation time step was taken to be *τ*_*s*_ = 0.008 *τ*_0_. The simulations were equilibrated for a time of at least 300 000 *τ*_*s*_ and further run for 5 000 000 *τ*_*s*_ (2 000 000 *τ*_*s*_ for the main Figs. 5a-b and for Fig. 6). In order to gather enough statistics, each set of simulations was run for a number of different initial velocity random seeds: 300 for Fig. 2, at least 200 for Fig. 3, 400 for the insets of Fig. 5a-b, 500 for Fig. 5c, and 100 for Fig. 6.

To compute potentials of mean force (main Figs. 5a-b), we used the Weighted Histogram Analysis Method [46] with a harmonic restraint of constant *k*_bias_ = 1 *k*_B_*T/σ*^2^, over 50 equally spaced windows, each of them being simulated for 20 different random seeds.

We used the LAMMPS molecular dynamics software to run the simulations [47, 48] and the OVITO software to visualise trajectory files [49].

### B. Theoretical model

The constrained free energy *F* (*A*), for a given value *A* of membrane surface adhered to the nanoparticle, can be obtained as the sum of the following terms.

The bending energy is

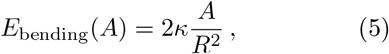

where *κ* is the bending rigidity, *R* is the radius of the nanoparticle plus the radius of a membrane bead, and the factor 2 accounts for both directions of curvature.

The adhesion energy is

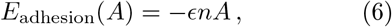

where *n* is the surface density of beads on the membrane, and *ϵ* is the average energy gain per bead when adhered to the nanoparticle surface.

The surface energy term is due to an adhesion-induced stretching of the membrane. It reads

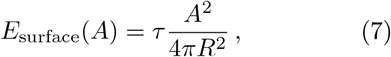

where *τ* is the surface tension. Adhesion can indeed stretch a membrane, shaping an unbound disk of membrane into a spherical cap, with an area increase that is quadratic in the bound area *A*. This term is effectively irrelevant in our simulations, that are performed at constant zero pressure.

The entropic contribution represents in a qualitative manner the decrease in entropy due to adhesion and consequent confinement of membrane beads close to the nanoparticle. It is computed as

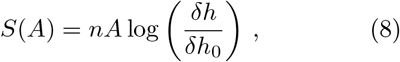

within a 1D-ideal-gas approximation for membrane beads, as far as motion perpendicular to the membrane surface is concerned. *δh* is the typical amplitude of fluctuations for a bound membrane surface *A*, while *δh*_0_ is the same quantity for a surface *A* of the reference (unbound) membrane. Lengths *δh* and *δh*_0_ are estimated in the following way. First we write down a constrained Helfrich Hamiltonian. For the bound state, this reads

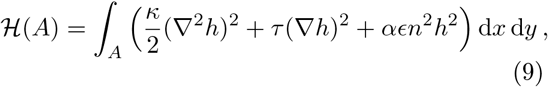

where *h* is the distance of the membrane at (*x, y*) from its ground-state surface *xy*. The first term in parentheses represents (microscopic) bending, the second surface tension, and the third is a quadratic expansion of the adhesion potential around its minimum, with *α* = 62 · 2^2*/*3^ ≃ 100 if the considered potential is a Lennard-Jones with energy parameter *E*. For the unbound reference state, the Hamiltonian reads the same, except for the last term which is obviously not present (*α* = 0). Writing *h* by means of a Fourier series allows us to compute thermal averages 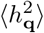 (or 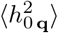 for the unbound state) through the equipartition theorem:

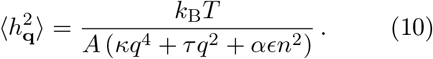

From here it is clear that the role of adhesion is to penalise small wavenumbers *q* = |**q**|, i.e., in a tensionless membrane (*τ* = 0), wavelengths shorter than 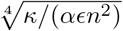. We then integrate 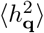 for wavelengths ranging from the size of a bead 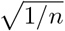 to the size of the adhered membrane portion 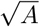: this integral provides an upper-bound estimate for the square of the typical fluctuation amplitude *δh* (or *δh*_0_), at a given wrapped area *A*. The result is

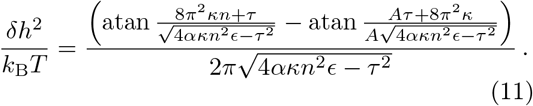

For *τ* = 0, this can be expanded as follows:

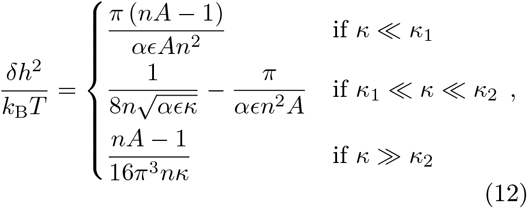

with

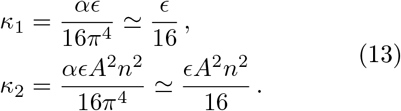

In practice: for *κ* ≪ *κ*_1_, all fluctuation modes are adhesion-dominated; for *κ*_1_ ≪ *κ* ≪ *κ*_2_, fluctuations are limited by adhesion on a wavelength 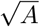, but by bending on a shorter wavelength 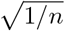: at larger *κ*, all permitted fluctuations are bending-dominated. For the reference unbound state, it is always [50, 51]

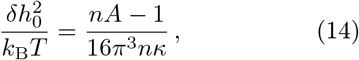

since *κ*_1_ and *κ*_2_ from Eq. (13) go to 0, as *α* goes to 0. Fluctuations can only be bending-dominated in this case.

Now, given Eqs. (12) and (14), the entropy of the adhered state with respect to the free one is provided by Eq. (8). A comparison between Eqs. (12) and (14) shows that the entropy gain is maximal, in absolute value, for small *κ*, and decreases to 0 as *κ* approaches *κ*_2_.

Fig. 4 is obtained by numerically minimising the total constrained free energy, resulting from the sum of (5), (6), (7) and (8), with respect to the bound surface *A*; then, the total free energy is computed, together with its bending, adhesion and entropic terms, for such optimal value of *A*. In the calculation, we use parameters meant to represent our molecular dynamics simulations: *n*^*−*1*/*2^ = 1.14 *σ* (where *σ* is the Lennard-Jones length unit), *R* = 6.17 *σ, E* = 0.7 *k*_B_*T*, and *τ* = 10^*−*3^ *k*_B_*T/σ*^2^.

## References

[1] Martin GL Van den Heuvel and Cees Dekker. Motor proteins at work for nanotechnology. Science, 317(5836):333–336, 2007.

[2] Ronald D Vale. The molecular motor toolbox for intracellular transport. Cell, 112(4):467–480, 2003.

[3] Joseph R Casey, Sergio Grinstein, and John Orlowski. Sensors and regulators of intracellular ph. Nature reviews Molecular cell biology, 11(1):50– 61, 2010.

[4] Amelia Barreiro, Riccardo Rurali, Eduardo R Hernández, Joel Moser, Thomas Pichler, Laszlo Forro, and Adrian Bachtold. Subnanometer motion of cargoes driven by thermal gradients along carbon nanotubes. Science, 320(5877):775–778, 2008.

[5] BC Regan, S Aloni, RO Ritchie, U Dahmen, and A Zettl. Carbon nanotubes as nanoscale mass conveyors. Nature, 428(6986):924–927, 2004.

[6] Philipp AE Schoen, Jens H Walther, Dimos Poulikakos, and Petros Koumoutsakos. Phonon assisted thermophoretic motion of gold nanoparticles inside carbon nanotubes. Applied Physics Letters, 90(25):253116, 2007.

[7] Mario Tagliazucchi and Igal Szleifer. Transport mechanisms in nanopores and nanochannels: can we mimic nature? Materials Today, 18(3):131– 142, 2015.

[8] Jan Genzer and Rajendra R. Bhat. Surface-Bound Soft Matter Gradients. Langmuir, 24(6):2294– 2317, 2008.

[9] Chun-Min Lo, Hong-Bei Wang, Micah Dembo, and Yu-li Wang. Cell movement is guided by the rigidity of the substrate. Biophysical journal, 79(1):144–152, 2000.

[10] Raimon Sunyer and Xavier Trepat. Durotaxis. Current Biology, 30(9):R383–R387, 2020.

[11] Adam Shellard and Roberto Mayor. Durotaxis: The Hard Path from In Vitro to In Vivo. Developmental Cell, 56(2):227–239, 2021.

[12] Charles R. Doering, Xiaoming Mao, and Leonard M. Sander. Random walker models for durotaxis. Physical Biology, 15(6):066009, 2018.

[13] Tienchong Chang, Hongwei Zhang, Zhengrong Guo, Xingming Guo, and Huajian Gao. Nanoscale directional motion towards regions of stiffness. Physical review letters, 114(1):015504, 2015.

[14] Chao Wang and Shaohua Chen. Motion driven by strain gradient fields. Scientific reports, 5:13675, 2015.

[15] Robert W Style, Yonglu Che, Su Ji Park, Byung Mook Weon, Jung Ho Je, Callen Hyland, Guy K German, Michael P Power, Larry A Wilen, John S Wettlaufer, et al. Patterning droplets with durotaxis. Proceedings of the National Academy of Sciences, 110(31):12541–12544, 2013.

[16] Jesus Bueno, Yuri Bazilevs, Ruben Juanes, and Hector Gomez. Wettability control of droplet durotaxis. Soft matter, 14(8):1417–1426, 2018.

[17] Panagiotis E Theodorakis, Sergei A Egorov, and Andrey Milchev. Stiffness-guided motion of a droplet on a solid substrate. The Journal of Chemical Physics, 146(24):244705, 2017.

[18] Donald M Engelman. Membranes are more mosaic than fluid. Nature, (438):578–580, 2005.

[19] Erdinc Sezgin, Ilya Levental, Satyajit Mayor, and Christian Eggeling. The mystery of membrane organization: Composition, regulation and roles of lipid rafts. Nature Reviews Molecular Cell Biology, 18(6):361–374, 2017.

[20] Milka Doktorova, Jessica L. Symons, and Ilya Levental. Structural and functional consequences of reversible lipid asymmetry in living membranes. Nature Chemical Biology, 16(12):1321–1330, 2020.

[21] Amirali Hossein and Markus Deserno. Spontaneous Curvature, Differential Stress, and Bending Modulus of Asymmetric Lipid Membranes. Biophysical Journal, 118(3):624–642, 2020.

[22] Alexandre Santinho, Aymeric Chorlay, Lionel Foret, and Abdou Rachid Thiam. Fat Inclusions Strongly Alter Membrane Mechanics. Biophysical Journal, 2021.

[23] Claire Valotteau, Andra C. Dumitru, Laurence Lecordier, David Alsteens, Etienne Pays, David Pérez-Morga, and Yves F. Dufrêne. Multiparametric Atomic Force Microscopy Identifies Multiple Structural and Physical Heterogeneities on the Surface of Trypanosoma brucei. ACS Omega, 5(33):20953–20959, 2020.

[24] Juan Mucci, Andrés B. Lantos, Carlos A. Buscaglia, María Susana Leguizamón, and Oscar Campetella. The Trypanosoma cruzi Surface, a Nanoscale Patchwork Quilt. Trends in Parasitology, 33(2):102–112, 2017.

[25] Yong Keun Park, Catherine A. Best, Kamran Badizadegan, Ramachandra R. Dasari, Michael S. Feld, Tatiana Kuriabova, Mark L. Henle, Alex J. Levine, and Gabriel Popescu. Measurement of red blood cell mechanics during morphological changes. Proceedings of the National Academy of Sciences of the United States of America, 107(15):6731–6736, 2010.

[26] Laura Picas, Félix Rico, Maxime Deforet, and Simon Scheuring. Structural and Mechanical Heterogeneity of the Erythrocyte Membrane Reveals Hallmarks of Membrane Stability. ACS Nano, 7(2):1054–1063, 2013.

[27] Adi de la Zerda, Michael J. Kratochvil, Nicholas A. Suhar, and Sarah C. Heilshorn. Review: Bioengineering strategies to probe T cell mechanobiology. APL Bioengineering, 2(2):021501, 2018.

[28] Charles Roduit, F. Gisou Van Der Goot, Paolo De Los Rios, Alexandre Yersin, Pascal Steiner, Giovanni Dietler, Stefan Catsicas, Frank Lafont, and Sandor Kasas. Elastic membrane heterogeneity of living cells revealed by stiff nanoscale membrane domains. Biophysical Journal, 94(4):1521–1532, 2008.

[29] Frederick A Heberle, Milka Doktorova, Haden L Scott, Allison D Skinkle, M Neal Waxham, and Ilya Levental. Direct label-free imaging of nanodomains in biomimetic and biological membranes by cryogenic electron microscopy. Proceedings of the National Academy of Sciences, 117(33):19943–19952, 2020.

[30] Philip W. Fowler, Jean Hélie, Anna Duncan, Matthieu Chavent, Heidi Koldsø, and Mark S. P. Sansom. Membrane stiffness is modified by integral membrane proteins. Soft Matter, 12(37):7792–7803, 2016.

[31] Nicola Mandriota, Claudia Friedsam, John A. Jones-Molina, Kathleen V. Tatem, Donald E. Ingber, and Ozgur Sahin. Cellular nanoscale stiffness patterns governed by intracellular forces. Nature Materials, 18(10):1071–1077, 2019.

[32] Anita Joanna Kosmalska, Laura Casares, Alberto Elosegui-Artola, Joseph Jose Thottacherry, Roberto Moreno-Vicente, Víctor González-Tarragó, Miguel Ángel Del Pozo, Satyajit Mayor, Marino Arroyo, Daniel Navajas, Xavier Trepat, Nils C. Gauthier, and Pere Roca-Cusachs. Physical principles of membrane remodelling during cell mechanoadaptation. Nature Communications, 6, 2015.

[33] Kai Yang, Ran Yang, Xiaodong Tian, Kejie He, Seth Leon Filbrun, Ning Fang, Yuqiang Ma, and Bing Yuan. Partitioning of nanoscale particles on a heterogeneous multicomponent lipid bilayer. Physical Chemistry Chemical Physics, 20(44):28241–28248, 2018.

[34] Chapin S Korosec, Lavisha Jindal, Mathew Schneider, Ignacio Calderon de la Barca, Martin J Zuckermann, Nancy R Forde, and Eldon Emberly. Substrate stiffness tunes the dynamics of polyvalent rolling motors. Soft Matter, 2021.

[35] Reinhard Lipowsky. The conformation of membranes. Nature, 349(6309):475–481, 1991.

[36] Reinhard Lipowsky. Generic interactions of flexible membranes, volume 1. Elsevier Masson SAS, 1995.

[37] Udo Seifert. Configurations of fluid membranes and vesicles. Advances in Physics, 46(1):13–137, 1997.

[38] Elizaveta A Novikova, Matthew Raab, Dennis E Discher, and Cornelis Storm. Persistence-driven durotaxis: generic, directed motility in rigidity gradients. Physical review letters, 118(7):078103, 2017.

[39] Ziyang Xu, Lijuan Gao, Pengyu Chen, and LiTang Yan. Diffusive transport of nanoscale objects through cell membranes: a computational perspective. Soft Matter, 16(16):3869–3881, 2020.

[40] Ellen Reister-Gottfried, Stefan M. Leitenberger, and Udo Seifert. Hybrid simulations of lateral diffusion in fluctuating membranes. Physical Review E - Statistical, Nonlinear, and Soft Matter Physics, 75(1):1–11, 2007.

[41] Ali Naji and Frank L.H. Brown. Diffusion on ruffled membrane surfaces. Journal of Chemical Physics, 126(23), 2007.

[42] M. E. Cates. Diffusive transport without detailed balance in motile bacteria: does microbiology need statistical physics? Reports on Progress in Physics, 75(4):042601, 2012.

[43] Kathryn A. Rosowski, Tianqi Sai, Estefania Vidal-Henriquez, David Zwicker, Robert W. Style, and Eric R. Dufresne. Elastic ripening and inhibition of liquid–liquid phase separation. Nature Physics, 16(4):422–425, 2020.

[44] Estefania Vidal-Henriquez and David Zwicker. Theory of droplet ripening in stiffness gradients. Soft Matter, 16(25):5898–5905, 2020.

[45] Stephen Sung and David Chandler. Perturbation theory for repulsive forces in classical fluids: selected applications. The Journal of Chemical Physics, 56(10):4989–4994, 1972.

[46] Shankar Kumar, John M Rosenberg, Djamal Bouzida, Robert H Swendsen, and Peter A Kollman. The weighted histogram analysis method for free-energy calculations on biomolecules. i. the method. Journal of computational chemistry, 13(8):1011–1021, 1992.

[47] Steve Plimpton. Fast parallel algorithms for short-range molecular dynamics. Journal of computational physics, 117(1):1–19, 1995.

[48] P. J. in ‘t Veld, S. J. Plimpton, and G. S. Grest. Accurate and efficient methods for modeling colloidal mixtures in an explicit solvent using molecular dynamics. Comp. Phys. Comm., 179:320– 329, 2008.

[49] Alexander Stukowski. Visualization and analysis of atomistic simulation data with ovito–the open visualization tool. Modelling and Simulation in Materials Science and Engineering, 18(1):015012, 2009.

[50] David Boal. Membrane undulations. In Mechanics of the Cell, volume 1, chapter 8, pages 292–325. Cambridge University Press, Cambridge, 2012.

[51] W. Helfrich and R. M. Servuss. Undulations, steric interaction and cohesion of fluid membranes. Il Nuovo Cimento D, 3(1):137–151, 1984.

[52] Tine Curk, Peter Wirnsberger, Jure Dobnikar, Daan Frenkel, and Anđela Šarić. Controlling Cargo Trafficking in Multicomponent Membranes. Nano Letters, 18(9):5350–5356, 2018.

[53] κ (or κ_soft_ or κ_stiff_) is the obvious counterpart of the numerical parameter K_b_ (or K_soft_ or K_stiff_). The exact mapping between κ in our theory and K_b_ in our simulations is not important here. For an example of such conversion, see [52].

